# Behavioral and neural evidence of the rewarding value of exercise behaviors: A systematic review

**DOI:** 10.1101/211425

**Authors:** Boris Cheval, Rémi Radel, Jason L. Neva, Lara A. Boyd, Stephan P. Swinnen, David Sander, Matthieu P. Boisgontier

**Affiliations:** Swiss NCCR “LIVES - Overcoming Vulnerability: Life Course Perspectives”, University of Geneva, Switzerland; Department of General Internal Medicine, Rehabilitation and Geriatrics, University of Geneva, Switzerland; Laboratoire Motricité Humaine Expertise Sport Santé (LAMHESS), Université Côte d’Azur, Nice, France; Brain Behavior Laboratory, University of British Columbia, Vancouver, British Columbia, Canada; Movement Control and Neuroplasticity Research Group, Department of Movement Sciences, Biomedical Sciences Group, KU Leuven, Leuven, Belgium; Leuven Research Institute for Neuroscience and Disease (LIND), KU Leuven, Leuven, Belgium; Swiss Center for Affective Sciences, University of Geneva, Geneva, Switzerland; Laboratory for the Study of Emotion Elicitation and Expression, Department of Psychology, University of Geneva, Geneva, Switzerland

## Abstract

**Background:** In a time of physical inactivity pandemic, attempts to better understand the factors underlying the regulation of exercise behavior are important. The dominant neurobiological approach to exercise behavior considers physical activity to be a reward. However, negative affective responses during exercise challenge this idea.

**Objective:** Our objective was to systematically review studies testing the automatic reactions triggered by stimuli associated with different types of exercise behavior (e.g., physical activity, sedentary behaviors) and energetic cost variations (e.g., decreased energetic cost, irrespective of the level of physical activity).

**Methods:** Two authors systematically searched, screened, extracted, and analyzed data from articles in the MEDLINE database.

**Results:** We included 26 studies. Three outcomes of automatic processes were tested: Affective reactions, attentional capture, and approach tendencies. Behavioral results show that physical activity can become attention-grabbing, automatically trigger positive affect, and elicit approach behaviors. These automatic reactions explain and predict exercise behaviors. However, the use of a wide variety of measures prevents drawing solid conclusions about the specific effects of automatic processes. Brain imaging results are scarce but show that stimuli associated with physical activity and, to a lesser extent, sedentary behaviors activate regions involved in reward processes. Studies investigating the rewarding value of behaviors driving energetic cost variations such as behaviors minimizing energetic cost are lacking.

**Conclusion:** Reward is an important factor in exercise behavior. The literature based on the investigation of automatic behaviors seems in line with the suggestion that physical activity is rewarding, at least for physically active individuals. Results suggest that sedentary behaviors could also be rewarding, although this evidence remains weak due to a lack of investigations. Finally, from an evolutionary perspective, behaviors minimizing energetic cost are likely to be rewarding. However, no study has investigated this hypothesis. In sum, additional studies are required to establish a strong and complete framework of the reward processes underlying automatic exercise behavior.

**Key points:**

- Behavioral and brain imaging studies using different outcomes of automatic behavior show that physical activity and, to a weaker extent, sedentary behaviors are rewarding.
- Behaviors minimizing energetic cost have been essential to evolutionary survival and are likely to be rewarding. However, experimental evidence is still lacking.
- The dominant neuropsychological approaches to exercise behavior are incomplete, which may partly explain our current inability to counteract the pandemic of physical inactivity.

## 1. Introduction

Twenty years ago, the World Health Organization (WHO) issued comprehensive guidelines for promoting physical activity among older adults [1]. Since then, the importance of physical activity for health has been increasingly emphasized and guidelines have been extended to all populations [2]. Today, however, one-third of the adult population remains physically inactive and 80% of the adolescent population does not reach the recommended amount of physical activity [3]. Why do most people fail to exercise regularly [4]? What if a fundamental principle that leads us to minimize energetic cost has been neglected in exercise neuropsychology?

In neurobiology, physical activity is considered to be a reward [5-10]. Yet, recent review [11], opinion [12], and conceptual articles [13] in the field of psychology underlined the displeasure experienced during exercise, which challenges the idea that physical activity is rewarding. So far, studies mainly focused on the automatic reactions triggered by physical activity. Studies investigating the automatic reactions triggered by sedentary behaviors remain scarce and no study has examined whether energetic cost variations (increase vs. decrease) could be rewarding irrespective of the considered level of physical activity.

Here, we examine evidence supporting the hypothesis that behaviors minimizing energetic cost (BMEC) are rewarding and, as such, are associated with a positive affective valence. This new concept based on the variation rather than the level of energetic cost could complete the current approach to exercise behavior and has the potential to enhance our understanding of the basic neurophysiological processes governing automatic processes in this field. In addition to providing new fundamental knowledge, this new approach could help to address a global health problem. Each year, physical inactivity is responsible for 13 million lost years of healthy life [14] and 5 million deaths worldwide [15]. To counteract the pandemic of physical inactivity [4], reconsidering the fundamental basis of the current approach to exercise behavior is needed.

In this introduction section, we first explain that the regulation of exercise behavior is based on two types of processes (controlled and automatic). Second, we define reward, highlight its brain substrates, and explain how it triggers automatic processes. Third, we demonstrate that BMEC could be rewarding. Fourth, we describe the evolutionary basis of the rewarding value of BMEC and discuss how, in occidental societies, this rewarding value could activate automatic processes increasing the inertia of physical inactivity. Fifth, we examine the implications of such a reward. Finally, we set the objectives of this systematic review on automatic exercise behavior.

### 1.1. Controlled and automatic processes in exercise behavior

In neuroscience and psychology, two types of processes are thought to govern the regulation of behaviors: Controlled and automatic processes [16-19]. Controlled processes are initiated intentionally, require cognitive resources, and operate within conscious awareness. Conversely, automatic processes are initiated unintentionally, tax cognitive resources to a much lesser extent, and occur outside conscious awareness [20,21]. These automatic processes can be problematic when they come into conflict with the controlled processes [19,22]. For example, an opportunity for sedentary behavior can automatically activate a behavioral response that competes with the conscious intention to adopt a physically active behavior, thereby preventing its implementation. This conflict was recently highlighted in the affective-reflective theory of physical activity and exercise. This theory states that inactive individuals can fail to implement their motivation to change from an inactive to an active state because of a restraining action impulse resulting from the pleasurable affect associated with being at rest [12]. Models testing the capacity of controlled processes for explaining exercise behavior have shown high levels of unexplained variance [23]. Concurrently, automatic processes may play an important role in the prediction of exercise behavior [24-26]. The pervasive effects of automatic on exercise behavior [27] suggest that the pandemic of physical inactivity [4] originates, at least in part, in automatic processes. Specifically, individuals may fail to exercise regularly despite conscious intentions to be active because BMEC activate competing automatic processes. Here, BMEC are defined as any behavior resulting in energetic cost decrease, irrespective of the initial level of energetic expenditure. Whether BMEC are automatically triggered due to a positive affective valence that is typical of a rewarding behavior is still unknown.

### 1.2. Reward, automatic processes, and brain substrates

Reward has been investigated using multiple techniques (e.g., neuroimaging, electrophysiology, pharmacology) and model organisms (e.g., rodent, zebrafish, monkeys) to understand different processes and states in humans (e.g., development, aging, obesity, addiction). Reward is the positive value ascribed to an object, a behavioral act, or an internal physical state [28], through multiple neuropsychological components [29-31]. The “wanting” (or desire) component is the positive value resulting from the relevance of the behavior for the needs of the individual [30, 32, 33]. The “liking” component is the positive value resulting from the hedonic pleasure associated with the performance of the behavior [29]. These components share a neural substrate in the ventral pallidum, amygdala, nucleus accumbens, and striatum (which includes the putamen, caudate, and globus pallidus). However, it has been suggested that the networks they rely on are not strictly identical. Wanting relies on the premotor cortex, central nucleus of the amygdala, nucleus accumbens core, putamen, caudate, and ventral pallidum [34-39], whereas liking relies on the nucleus accumbens shell, ventral pallidum, orbitofrontal cortex, insular cortex, and parabrachial nucleus [33,40-47]. In humans, most of the studies investigating the neural substrates of reward relied on food or addictive substances, such as cocaine, alcohol, or nicotine [48]. Based on these studies, systematic reviews have been conducted [49-51] and showed that food and addictive cues activate the brain regions associated with reward. However, no literature synthesis has been undertaken investigating the rewarding value of physical activity, sedentary behaviors, or BMEC.

Because a reward has a positive value, individuals are inclined to obtain it. In this perspective, a reward has an incentive function. The perception of reward-related cues triggers the release of dopamine in the brain, which leads these cues to become attention-grabbing, induce subjective craving, and ultimately elicit approach behavior. These processes are generally assumed to be automatic as they have been shown using speeded reaction-time tasks [52,53], pupil response [54-55], and early event-related potential [56-57]. These automatic processes have presumably evolved to speed up the processing of threats and rewards in the environment. This faster processing confers an advantage for survival [58]. It should be noted that the term reward may have different meanings in neuroscience, biology, and psychology. In this study, reward refers to stimuli with an incentive value because of the positive affective hedonic experiences previously associated with them.

### 1.3. Behaviors minimizing energetic cost as a reward

To counteract the lack of physical activity, reconsidering our current view of the psychological and neural mechanisms regulating exercise behavior is urgently needed. In this study, we contend that a fundamental principle pushing individuals to minimize energetic cost has been insufficiently considered in the dominant approaches to exercise behavior. Reward triggers automatic processes that can initiate, sustain, and change behavior adaptively between different available options and plays a key role in optimizing the allocation of resources necessary for evolutionary survival [59]. BMEC determine behavioral adaptation on short (e.g., walking vs. running [60]) and evolutionary timescales (e.g., quadrupedalism vs. bipedalism [61]). Therefore, BMEC could be conceived of as a reward. This assumption concurs with previous studies claiming that individuals possess an innate tendency to conserve energy and avoid unnecessary physical exertion [62,63], thereby explaining the negative affects experienced during exercise [13]. This evolutionary view of exercise may explain the exercise paradox: Why do individuals persist to be physically inactive despite knowledge of the risks associated with this inactivity? In the following section, we elaborate on the suggested evolutionary explanation of this paradox.

### 1.4. From an automatic to a controlled trigger of physical activity

Besides BMEC, physical activity has been necessary for development and evolutionary survival. For example, physical activity can be viewed as a necessary means to achieve motor learning and development, explaining why children are naturally inclined to exert physical effort in play periods [64]. Furthermore, physical activity is triggered when individuals need to search for food or shelter, interact with competitors, and avoid predators [65]. Particularly, food and physical activity are thought to be part of the same cycle, where alternating periods of food scarcity and abundance are associated with higher and lower physical activity, respectively [66]. This increase of physical activity during food restriction is interpreted as foraging behavior that conferred a decisive advantage for survival in periods of food scarcity [67-69]. Since the goal of this increased energetic expenditure is energy replenishment, the optimization principle is at work. This principle is based on cost minimization [70]. For example, individuals automatically adapt their step frequency and walking speed in real-time to optimize energetic costs [71], and learn to minimize the physical efforts required to obtain a specific reward [72]. In addition, individuals who sustained physical activity for longer periods were more likely to find food. Therefore, species that developed processes alleviating pain and fatigue during physical activity, such as analgesia, sedation, and anxiolysis [5-10], were more likely to survive. Physical activity likely became a reward due to these convergent processes.

In modern occidental societies, the food-physical activity circle is broken. Food scarcity, the automatic trigger of physical activity, no longer exists. Individuals engage in physical activity because of controlled processes, such as conscious intentions to regulate energetic balance. When engaged in regular physical activity and exercise, the repetition of hedonic experiences can be learned [73] and result in the association of the hedonic effect and the behavior (e.g., running). Once this automatic association is consolidated in memory, an environmental stimulus (e.g., seeing another individual running) can automatically trigger a positive evaluation of the stimulus, which will, in turn, evoke preparatory responses favoring the engagement in running, such as approach tendencies toward running. This rationale is in line with the affective-reflective theory of physical inactivity and exercise and a growing number of studies showing that the positive affective valence of exercise is associated with subsequent exercise [13]. Additionally, the ability to develop automatic strategies counteracting the rewarding value of sedentary behaviors has been suggested in a recent study showing that sedentary behaviors primes facilitated the recognition of words associated with physical activity, only in individual who manage to maintain regular physical activity [74].

### 1.5. Implications

In the fields of psychology and neuroscience, the neutral nature of sitting and lying positions has always been taken for granted. If BMEC are a reward, this assumption suddenly becomes questionable. Reward perception has been shown to be dependent on individuals’ physiological state [75]. For example, thirsty participants show higher perceptual readiness to drinking-related stimuli [76] and hungry participants show stronger automatic approach reactions toward food-related stimuli [77]. Therefore, if BMEC are a reward, the reward associated with behaviors involving low levels of energetic costs (e.g., sitting and lying positions) or minimizing energetic expenditure (e.g., taking the escalator or elevator instead using of the stairs) depends on multiple factors such as participant’s maximal exercise capacity and recent exercise history (e.g., did the participant come by bike or bus). Accordingly, it is urgent to investigate the rewarding nature of BMEC. Until this point has been clarified, a precautionary principle should be applied and scientists in the field of neuroscience, psychology, and exercise should prospectively adjust their experimental designs to discard this potential bias. Physical activity should be monitored during the days/hours preceding the experiment and this information should be included in the models as a covariate.

Additionally, if BMEC are a reward, the pandemic of physical inactivity is driven by an automatic resistance to the intended engagement in exercise. Therefore, public health policies take the wrong approach. Part of the massive investment aiming at increasing conscious intentions to be active should be redirected toward the development of research projects aiming at understanding the mechanisms underlying this automatic resistance and interventions aiming at reducing it.

### 1.6. Objective

Rewarding stimuli trigger different outcomes of automatic processes such as attention capture (i.e., reward captures attention), affective reactions (i.e., reward produces hedonic pleasure), and approach tendencies (i.e., reward predisposes to physical approach). Here, our objective was to systematically review studies investigating automatic reactions to stimuli associated with different types of exercise behavior (e.g., physical activity and sedentary behaviors) and energetic cost variations (e.g., BMEC). Effect sizes were extracted to evaluate the magnitude of the association between automatic processes and exercise behaviors.

## 2. Methods

### 2.1. Search strategy

Figure 1 presents the flow diagram of our search strategy, which was performed by two investigators (MPB and BC). The potential studies were identified by searching the electronic MEDLINE database via PubMed. We searched for all available records starting from January 2000 until June 2017 using the following combination of keywords in the title or abstract of the article: (exercise OR “physical activity” OR “sedentary behavior”) AND (reward OR automatic OR impulsive OR implicit OR non-conscious). This review was conducted according to the Preferred Reporting Systematic reviews and Meta-Analyses (PRISMA) guidelines [78] but our protocol was not pre-registered.

**Figure 1.**
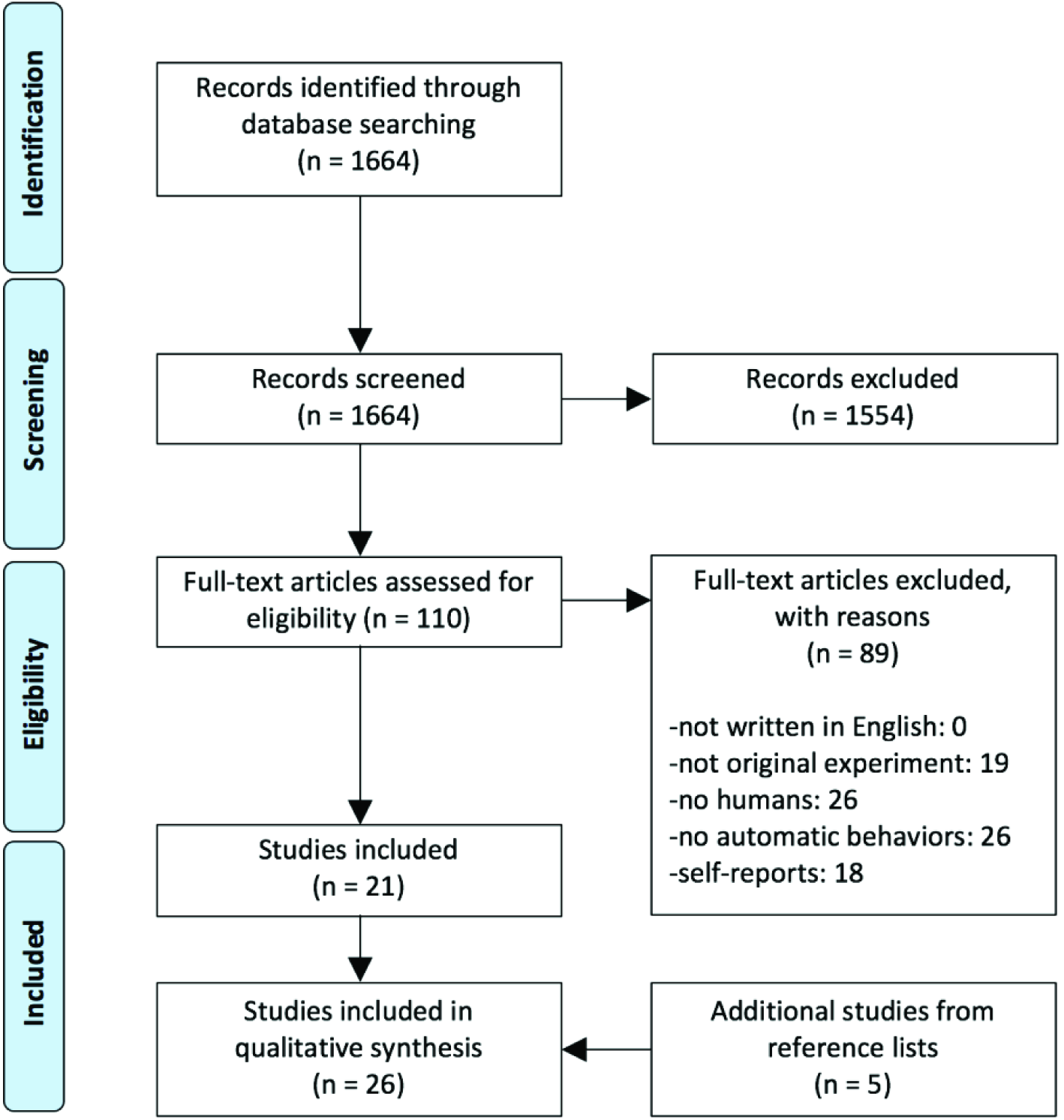
PRISMA flow diagram

### 2.2. Eligibility criteria and study selection

#### 2.2.1. Inclusion criteria

To be included in this systematic review, the article had to 1) be published in a peer-reviewed journal and written in English, 2) report original data collected from humans, 3) test the automatic behaviors or brain activation triggered by stimuli associated with different types of exercise behavior (e.g., physical activity, sedentary behaviors, and BMEC), and 4) assess these automatic behaviors and brain activation during the presentation of stimuli related to exercise behavior or during the performance of this behavior.

#### 2.2.2. Exclusion criterion

Self-reported measures were excluded as they appear to be less appropriate to measure the automatic processes associated with reward [79]. For a review of the relationship between self-reported habit index and exercise behavior, please see Gardner et al. [80].

#### 2.2.3. Exclusion selection

Five steps were used to select the articles meeting the inclusion criteria. If there was a doubt at any step, the article was kept for further inspection. At step 1, articles not written in English were excluded. At Step 2, articles not reporting original experimental data were excluded (e.g., reviews, meta-analyses, commentary, technical reports, case studies). At Step 3, articles were excluded if they did not involve a human population. At step 4, articles were excluded if they did not test the automatic behaviors or brain activation triggered by rewarding stimuli. At step 5, articles exclusively using self-reported measures of these processes were excluded. We performed reference screening and forward citation tracking on the articles remaining after step 5.

### 2.3. Data extraction

Data were extracted from the included articles and summarized in Table 1. In this table, we report 1) the type of population (e.g., age, healthy individuals, individuals with pathologies such as anorexia nervosa, respiratory disease, or obesity), 2) the technique used to investigate brain substrates (functional magnetic resonance imaging; fMRI), 3) the type of measure used to assess behavioral performance (e.g., reaction time), 4) the type of task used (i.e., Implicit Association Test, Manikin task, Visual Dot Probe Task, imagined scenarios), 5) the type of reward used and the format in which the reward was presented (pictures, words, imagined stimuli), 6) the content of the reward, i.e., whether the reward was related to specific sports and fitness (e.g., scheduled physical activity such as running, swimming) and sedentary-related activities (e.g., specific activity associated with a low energetic expenditure such as watching TV, playing video games) or more to the general concept of action/effort (e.g., active, energetic, vigorous) and inaction/rest (e.g., weak, frail, inactive), 7) whether recent physical activity history was controlled, and 8) the effect sizes, which were extracted and transformed into Cohen’s d when necessary [81].

**Table 1.**
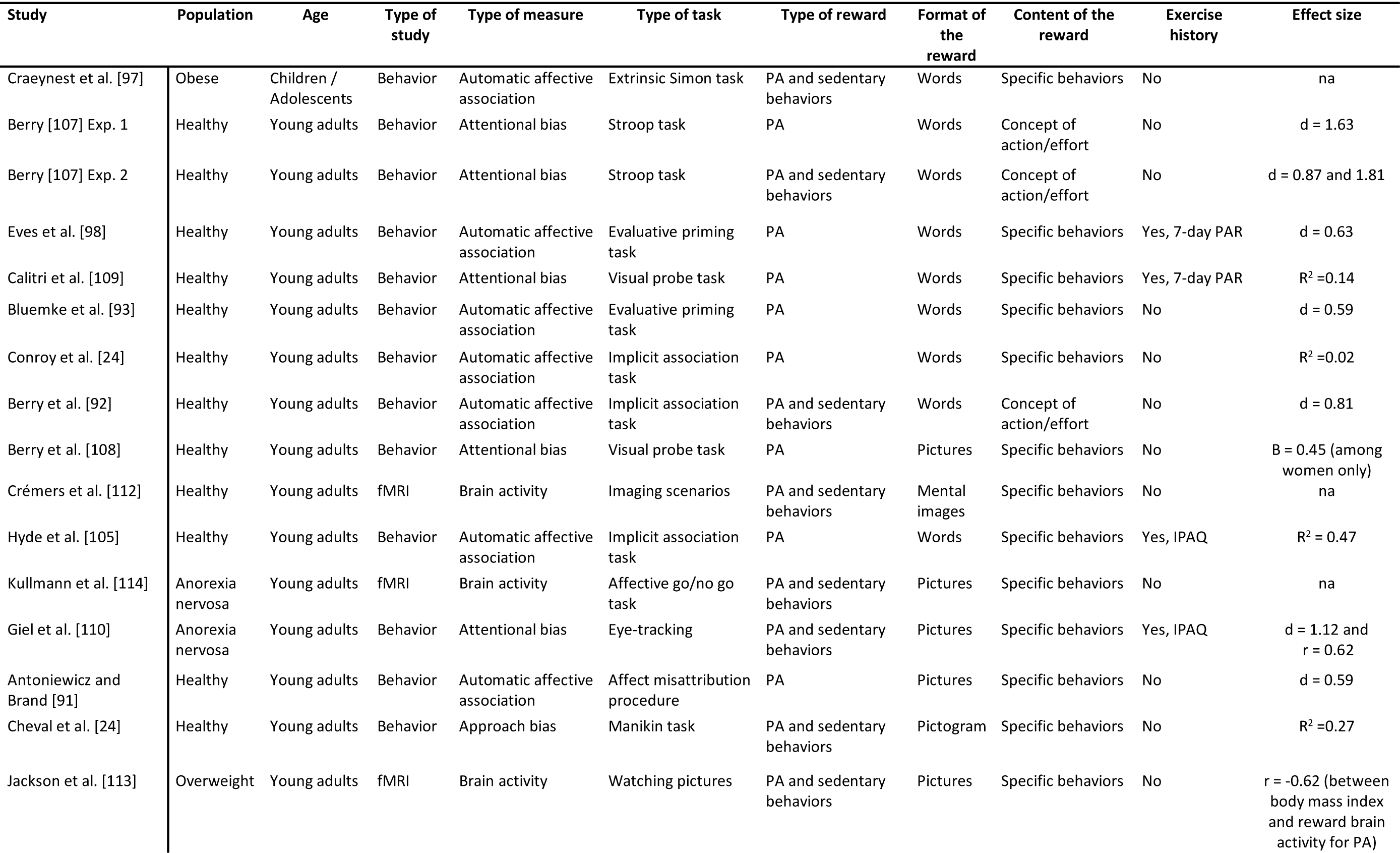
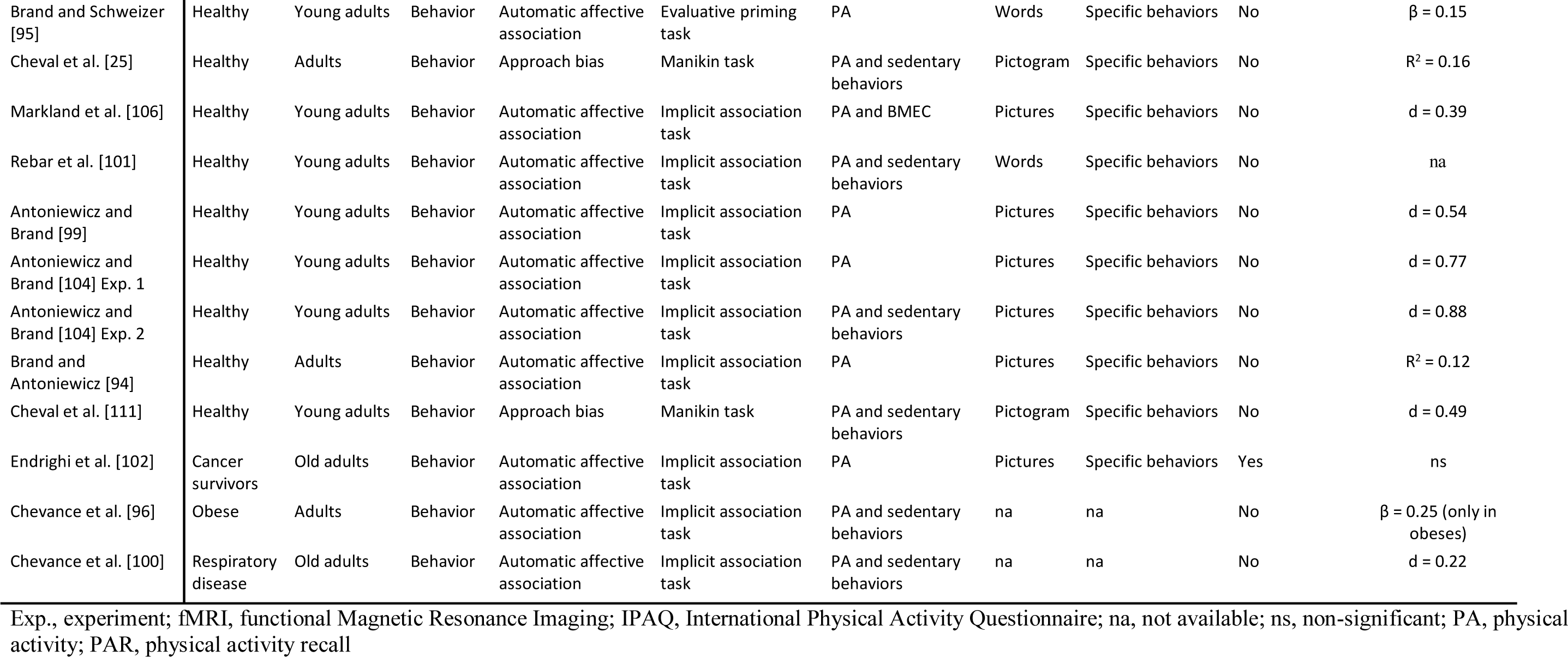
Studies investigating the automatic reactions and brain activations triggered by stimuli associated with physical activity and sedentary behaviors.

## 3. Results

### 3.1. Literature search

The primary search retrieved 1664 potentially relevant articles. Of the 1664 screened articles, a disagreement occurred in 31 cases (2%), which were all resolved by discussion. This selection yielded 110 potentially relevant full-text articles, which were then reviewed. All articles remained after step 1. At Step 2, 19 articles were excluded because they did not report original data. At Step 3, 26 articles were excluded because they did not involve humans. At step 4, 26 articles were excluded because they did not assess automatic reactions (e.g., affective reactions, attentional capture, and approach tendencies) triggered by stimuli associated with different types of exercise behaviors (e.g., physical activity, sedentary behaviors, and BMEC). At step 5, 18 articles were removed because they relied exclusively on self-reports. Five articles were added after reference screening and forward citation tracking. Finally, 26 articles were included (Table 1). Three of these 26 articles reported fMRI data.

### 3.2. Study characteristics

#### 3.2.1. Participants

Among the 26 included studies, 73.1% investigated healthy humans. The studies also investigated populations with weight control difficulties, such as overweight/obese individuals (11.6%) and anorexia nervosa patients (7.7%). A small set of studies investigated patients with a respiratory pathology (3.8%) or cancer survivors (3.8%) (Table 1). The studies investigated children (i.e., <18 years; 3.8%), young adults (i.e., > 19 and < 30 years; 77.0%), middle-aged adults (i.e., > 31 and < 49 years; 11.5%) and older adults (i.e., > 50 years; 7.7%) (Table 1).

#### 3.2.2. Tasks

Purely behavioral studies only using behavioral tasks represented 88.5% of studies and 11.5% also used fMRI. The studies investigated automatic affective associations (61.5%), attentional bias (15.4%), and approach tendencies (11.5%). Studies investigating automatic affective responses relied on the Implicit Association Test (IAT; 68.8%) [82], Evaluative Priming Task (18.8%) [83], Extrinsic Affective Simon Task (6.2%) [84], and Affect Misattribution Procedure (6.2%) [85]. Studies investigating attentional bias relied on the Visual Dot-Probe Task (VDP; 50.0%) [86], emotional Stroop Task (25.0%) [87], and eye-tracking (25.0%). Studies investigating approach tendencies relied on the manikin task [88,89]. Studies investigating the brain correlates mainly relied on imagined (33.3%) or watched (33.3%) scenarios associated with physical activity or inactivity, and on the go/no-go task (33.3%) [90,91].

#### 3.2.3. Reward

Studies investigated physical activity (46.2%) or both physical activity and sedentary behaviors (53.8%). None of the studies investigated BMEC. Words (38.5%), pictures (38.5%), pictograms (11.5%), and mental imagery (3.8%) were used as stimuli. Some studies did not explicitly indicate the format of the stimuli (7.7%). Most studies used stimuli associated with specific types of physical activity (e.g., tennis, football, swimming, walking) and sedentary behaviors (e.g., watching TV, reading a book, sitting in front of a computer; 84.6%). Some studies (7.7%) focused on the concept of action or effort (e.g., words like “active”, “energetic”, “vigorous”) and inaction or rest (e.g., words like “inactive”, “lethargic”, “lazy”), while the remaining 7.7% did not report the stimuli used.

#### 3.2.4. Recent exercise history

Recent exercise history was not controlled in 80.0% of the studies. The studies controlling for this state (20.0%) used a self-reported questionnaire (7-day physical activity recall or an adapted version of the International Physical Activity Questionnaire) to assess the amount of overall physical activity performed during the past week. One study used an accelerometer to measure this information.

### 3.3. Behavioral results

#### 3.3.1. Automatic affective processes

Sixteen studies were designed to investigate automatic affective processes. Six studies examined the differences in automatic affective reactions to stimuli associated with physical activity between physically active and inactive individuals [92-97]. Five of these studies showed the expected associations between automatic affective reactions and physical activity, but the effects sizes were highly variables (see table 1). For example, Bluemke et al. [94] used a priming task in which exercise (e.g., to swim, to jog) or control (e.g., to read, to eat) verbs were presented before participants had to quickly categorize positive (e.g., athletic, strong) and negative (e.g., exhausted, tense) target words. Results showed that physically active individuals were faster at categorizing positive target words after exercise primers, whereas inactive students were faster with negative words (d = 0.59). Using an IAT contrasting words associated with physical activity (e.g., workout, cross-train, run) and sedentary behaviors (sit, rest, snooze), a study revealed that individuals who were explicitly identified as exercisers had more positive automatic affective reactions toward exercise, as compared to non-exercisers (d = 0.81) [93]. Additionally, participants who reported greater habitual levels of physical activity also had more positive automatic affective reactions toward exercise compared to participants who reported less habitual physical activity levels. However, two studies did not show significant associations between automatic affective reactions and the level of physical activity [97] and weight status [98]. Specifically, using an Extrinsic Affective Simon Task with low (reading, resting, and watching television), moderate (walking, cycling, swimming), and high-intensity activities words (running, training, and exercising), results showed no differences in the automatic attitudes toward physical activity between a group of children with obesity and a matched control group [98]. Moreover, in an IAT task, results showed no significant associations between automatic affective reactions and the level of physical activity in the overall sample, but only among obese individuals (β = 0.25) [97].

Six studies also examined whether automatic affective reactions toward physical activity prospectively predict physical activity [26,99-103]. Five of these studies showed that positive automatic affective reactions toward physical activity prospectively predict physical activity [26,99-102], but the effects sizes were highly variables (see table 1). Using the Single Category Implicit Association Test (SC-IAT) [104], a variant of the IAT enabling the measurement of attitudes toward a specific target concept (e.g., physical activity only) rather than relative attitudes between two targets (physical activity vs. sedentary behaviors), Conroy et al. [26] showed that automatic affective reactions toward physical activity positively predicted the number of daily steps over one week, above and beyond controlled processes (e.g., behavioral intentions, outcome expectations). Although the association was statistically significant, automatic affective reactions accounted for only 2% of the additional variance in physical activity. Using the same SC-IAT, Rebar et al. [102] revealed that automatic affective reactions toward physical activity prospectively predicted the objectively measured level of physical activity over the next two weeks, above and beyond physical activity intentions. Furthermore, using an IAT contrasting stimuli associated with physical activity and sedentary behaviors, automatic affective reactions toward physical activity predicted adherence to a 14-week health and exercise course (d = 0.54) [99] and self-reported recreational physical activity six months after the end of a pulmonary rehabilitation program (d = 0.22) [100]. However, still using an IAT, a longitudinal study in endometrial cancer survivors did not demonstrate evidence supporting the fact that automatic affective reactions prospectively predicted daily minutes of exercise [103].

Finally, two studies examined how changes in automatic affective reactions were linked to physical activity [105,106]. The first study used the same SC-IAT and revealed that positive changes in affective reactions toward physical activity (i.e., from unfavorable to more favorable automatic evaluations) were associated with an increased self-reported physical activity over one-week (R^2^ = 0.47) [106]. The second study was designed to experimentally manipulate automatic affective reactions using an evaluative conditioning procedure [105]. Participants learned to associated pictures related to exercise (individuals engaging in individual and team sports, such as swimming or basketball) and non-exercise activities (individuals engaging non-physical activity such as watching TV or playing on a gaming console) with pictures associated with positive (individual relaxing in the sun) and negative (individual experiencing neck pain) affective feeling or experiences. Results revealed that participants who learned to associate exercise-related pictures with positive affective pictures and non-exercise-related pictures with negative affective pictures (i.e., acquisition of positive associations) reduced their negative automatic affective reactions toward physical activity (study 1, d = 0.77) and selected higher intensities on a self-paced cycling task compared to participants in a control condition (study 2, d = 0.88). A study showed that imagining a positive experience associated with physical activity led to more positive automatic affective reactions toward physical activity (d = 0.39) [107]. This study also showed more positive affective reactions toward physical activity in frequent exercisers (d = 0.57) [107].

In sum, most of the studies reported positive automatic affective reactions toward stimuli associated with physical activity. However, it is important to note that this positive association seems to apply to physical exercisers only. These results are in line with the rewarding value ascribed to physical activity. The positive bias reported on the affective reactions to stimuli associated with physical activity may reflect the “liking” component of the reward (hedonic pleasure).

#### 3.3.2. Attentional bias

Four studies were designed to investigate attentional bias [108-111]. Overall, physically active individuals showed an attentional bias toward stimuli associated with physical activity, whereas physically inactive individuals showed an attentional bias toward stimuli associated with sedentary behaviors [108-111]. One study used a Stroop color-naming task in which participants were instructed to quickly indicate the font color of words related to physical activity (e.g., energetic, vigorous, muscle), sedentary (e.g., unmotivated, lethargic, unfit), or control words (e.g., synthetic, suburban, varied) [108]. In this task, the difference in reaction time between exercise and control words and between sedentary and control words was used to infer the degree of attentional bias toward exercise and sedentary behaviors, respectively. Results revealed that regular exercisers (i.e., participants with an athletic identity) showed an attentional bias for exercise-related stimuli (study 1, d = 1.63; study 2, d = 0.87), whereas non-exercisers showed an attentional bias for sedentary-related words (study 2, d = 1.81). Another study using a VDP based on pairs of words, with one word related to physical activity associated with a neutral word (e.g., throw-cloth, football-sentence, tennis-devote), revealed a positive correlation between physical activity during the previous week and attentional bias toward words related to physical activity (β = 0.25) [110]. Another study used a VDP based on pairs of pictures, with one picture of an object related to exercise (e.g., football, stretching bands, field hockey stick, Frisbee) associated with a control picture where the exercise-related object was replaced by a non-exercise-related object (e.g., remote control, vacuum cleaner, beer bottle) [109]. Results showed an attentional bias toward physical activity in men, irrespective of their habitual level of physical activity, whereas only active women demonstrated such a bias toward physical activity (β = 0.45) [109]. Finally, a study tested attentional bias in adult patients with anorexia nervosa using eye-tracking [111]. Specifically, this study used a viewing task in which anorexia nervosa patients, physically active participants (i.e., at least 5 h per week of endurance sports), and physically inactive participants (i.e., only performing recreational physical exercise) were presented pairs of pictures, one related to an active situation (i.e., a young female athlete engaging in various physical activity) and one related to an inactive situation (i.e., a young female athlete engaging in various passive situations). They were instructed to freely explore picture pairs presented for 3 s on a computer screen. Results revealed that anorexia nervosa patients and physically active participants had a greater attentional bias toward stimuli associated with physical activity than physically inactive participants (d = 1.12). Additionally, in anorexia nervosa patients, attentional bias toward physical activity-related stimuli strongly correlated with the self-reported amount of physical activity (r = 0.62).

In sum, these results suggest that stimuli associated with physical activity can be attention-grabbing. However, it seems that this effect is specific to physical exercisers, although it could be extended to non-exercise men and anorexia nervosa patients. A study showed that sedentary stimuli could also be attention-grabbing.

#### 3.3.3. Automatic approach tendencies

Three studies were designed to investigate automatic approach tendencies toward physical activity and sedentary behaviors [24,25,112]. Overall, results showed that automatic approach tendencies toward physical activity positively predicted physical activity, whereas automatic approach tendencies toward sedentary behaviors negatively predicted physical activity [24,25]. All these studies used a manikin task based on pictograms representing physical activity and an active lifestyle (e.g., a pictogram of running, swimming, cycling) or rest and sedentary lifestyle (e.g., a pictogram of watching TV, lying on the sofa, resting). The first study showed that automatic approach tendencies toward physical activity predicted involvement in non-volitional physical activity in a laboratory context over and above intentions to be physically active, whereas automatic approach tendencies toward sedentary behaviors predicted lower involvement (R^2^ = 0.16) [24]. The second study extended these results in a more ecological context and revealed that these automatic approach tendencies predicted free time spent in physical activity over one week as measured with an accelerometer (R^2^ = 0.27) [25]. Finally, a study was designed to test whether automatic approach tendencies toward physical activity and sedentary behaviors can be manipulated using approach bias modification training, and subsequently impact exercise behaviors [112].

Results showed that participants trained to systematically approach physical activity and avoid sedentary behaviors spent longer periods of time exercising in the laboratory after the training compared to participants systematically trained to approach sedentary behaviors and avoid physical activity (d = 0.49) [112]. In sum, automatic approach tendencies toward stimuli depicting exercise behaviors, but also toward stimuli depicting sedentary behaviors, could be used to predict exercise behavior. The approach tendencies toward stimuli associated to exercise behaviors may reflect the “wanting” component of the reward (desire).

### 3.4. Brain substrates

Three studies reported potential brain substrates of reactions triggered by stimuli associated with physical activity and sedentary behaviors using fMRI. Results showed that some brain areas activated in response to these stimuli were consistent with areas highlighted in the reward literature [113-115]. However, an area shown to be involved in both the wanting and liking components of reward, the nucleus accumbens, was not reported in any of the 3 studies, thereby calling for further investigation. One study was conducted to identify the neural correlates involved in the control of brisk walking [113]. Young healthy individuals were asked to imagine themselves in three situations: Brisk walking in a long corridor, standing, and lying while their brain activity was measured using fMRI. Results revealed a stronger activation during mental imagery of brisk walking compared to mental imagery of standing or lying in areas associated with reward: Insula, pallidum and caudate. Another study examined inhibition response to active-and inactive-related stimuli [115] in anorexia nervosa patients, physically active participants (at least 5 h per week of endurance sports), and physically inactive participants (casual physical exercise) using a go/no-go task including stimuli associated with physical activity (e.g., a physically active person) and physical inactivity-(e.g., a physically inactive person) related pictures. The brain areas activated in this study were not related to reward. The last study tested the neural responses to pictures of physical activity and sedentary behaviors in a sample of overweight versus lean women [114]. Participants were asked to watch physical activities, sedentary activities, and landscape pictures presented during fMRI scanning. Results revealed an increased activation in brain areas associated with reward (amygdala, putamen, limbic lobe) when viewing pictures of physical activities compared to sedentary activities and control stimuli. Sedentary stimuli also activated the amygdala. Additionally, as body mass index increased, the activation of the right putamen decreased (r = −0.62). Finally, overweight women showed a decreased activation when watching sedentary compared with control stimuli in the insular cortex. These imaging results should be considered with caution as the experiments were not designed to investigate the brain regions associated with reward.

## 4. Discussion

The objective of this work was to systematically review studies investigating automatic reactions to stimuli associated with different types of exercise behavior (e.g., physical activity and sedentary behaviors) and energetic cost variations (e.g., BMEC). We included 26 studies and three outcomes of automatic processes were tested (affective reactions, attentional capture, and approach tendencies).

### 4.1. Main findings

Overall, results showed that automatic processes associated with physical activity are important for the regulation of physical activity. However, the use of a wide variety of measures, procedures, and targeted outcomes prevents strong conclusions, especially with regard to the effect sizes, which were highly variable [116,117]. Moreover, studies mainly rely on correlational designs, which provide a relatively low level of evidence. The few studies testing brain substrates of reward in exercise behavior showed that stimuli associated with physical activity increased activation in brain regions involved in reward (basal ganglia, amygdala, and prefrontal cortex). Most studies exclusively focused on stimuli depicting physical activity. Studies investigating sedentary behaviors were scarce and stimuli associated with energetic cost variations have not been investigated so far. These findings demonstrate that additional research investigating automatic reactions and brain substrates triggered by rewarding stimuli associated with physical activity, sedentary behaviors, and BMEC is required.

### 4.2. Current issues and perspectives

Besides the major findings reported above, this systematic review highlighted several theoretical and methodological issues that should be addressed in the future.

### 4.2.1. BMEC as a reward

The studies included in this review focused on automatic behaviors triggered by stimuli associated with physical activity. When stimuli associated with sedentary behavior were used, they were considered as a control condition most of the time. None of the studies were specifically designed to investigate the automatic behaviors or brain activation triggered by sedentary behaviors or BMEC. These findings reveal a knowledge gap in the literature of exercise behavior and highlight the necessity to address the potential rewarding value of BMEC and sedentary behaviors in future studies.

### 4.2.2. Neurophysiological studies are needed

This review showed a higher tendency to approach rather than avoid physical activity, irrespective of the individuals’ level of exercise. While this result seems in line with the suggestion that physical activity is a reward, it does not discard the possibility that sedentary behaviors and cost minimization are also rewarding [62]. First, almost all studies relied on physically active (motivated) individuals, who are more likely to have repeatedly experienced positive affective experiences during exercise than physically inactive (unmotivated) individuals [13]. As such, the results reported in the literature may be biased toward a higher rewarding value of physical activity. Second, this tendency to approach rather than avoid physical activity does not explain the fact that most individuals fail to be active despite their conscious intentions to be so. Yet, because all studies investigated behavioral outcomes (i.e., differences in reaction times), they may not provide a complete picture of the neural mechanisms underlying the automatic reactions in exercise behavior. For example, what we observe may result from facilitation processes but also from the competition of facilitation and inhibition processes supporting two different rewards (i.e., physical activity and BMEC). Purely behavioral work may not fully resolve this uncertainty and thus a neural approach is needed.

### 4.2.3. Recent exercise history

Very few studies controlled for exercise history over the days or hours preceding the experiment. This lack of control is an issue worth considering because, in other contexts, the perception of reward has been found to be dependent on the physiological state [75,118]. For instance, thirsty participants showed higher perceptual readiness to drinking-related stimuli [76] and hungry participants showed stronger automatic approach reactions toward food-related stimuli [34,119]. As mentioned in the Implications section, if BMEC are a reward, the reward associated with sitting and lying positions depends on the participant’s maximal exercise capacity and recent exercise history (e.g., did the participant come by bike or bus). Therefore, these factors should be considered during the conception of the study design.

### 4.2.4. Specific behaviors

This systematic review revealed that all the included studies focused on specific exercise behaviors (e.g., running, dancing, swimming) and/or sedentary behaviors (e.g., watching television, reading, video gaming). However, specific exercise behaviors can be rewarding due to the energetic cost they are associated with, but they can also be rewarding due to other factors. For example, the pleasure associated with a picture of an individual playing soccer may reflect the pleasure felt when watching this sport on TV, not the actual experience of playing soccer. The pleasure associated with a video gaming stimulus unlikely solely stems from the fact that the individual plays in a seated position. The pleasure associated with a skiing-related stimulus, the positive value ascribed to the stimulus (i.e., reward), may result from the liking component of speed perception, not only to value ascribed to the energetic cost associated with skiing. Therefore, it is difficult to infer strong conclusions from studies using an approach based on specific exercise behaviors.

## 5. Conclusion

Overall, results showed that automatic reactions toward stimuli depicting exercise behavior explained levels of physical activity. Imaging results showed that some brain regions associated with reward were activated by stimuli associated with physical activity and sedentary behaviors, but these studies remain scarce. These results highlight the importance of reward in exercise behavior.

This systematic review also reveals a knowledge gap that further highlights the necessity to reassess the veracity of the dominant neuropsychological approaches to exercise behavior. Neurophysiological techniques may afford the establishment of a strong and complete framework of reward in exercise behavior. Finally, this review illustrates an emerging line of research that has the potential to initiate the development of individualized and efficient interventions to counteract the pandemic of physical inactivity. For example, ensuring positive affective experiences during physical activity and exercise should result in automatic reactions favoring the engagement in physical activity [120]. Moreover, altering our environment in ways that reduce the opportunities to minimize energetic expenditure in favor of alternatives requiring higher intensities of physical activity could be particularly effective. Finally, interventions directly targeting automatic processes, such as evaluative conditioning and retraining automatic approach tendencies [22], could be effective in reducing the automatic attraction exerted by sedentary behaviors and BMEC.

## Compliance with Ethical Standards

### Funding

Matthieu Boisgontier is supported by a research grant (1501018N), a post-doctoral fellowship, and a grant for a long stay abroad from the Research Foundation—Flanders (FWO). The other authors report no sources of funding used to assist in the preparation of this article.

### Conflict of Interest

Boris Cheval, Rémi Radel, Jason Neva, Lara Boyd, Stephan Swinnen, David Sander, and Matthieu Boisgontier declare that they have no conflicts of interest relevant to the content of this review.

### Author contributions

Matthieu Boisgontier and Boris Cheval conceived the new model of exercise behavior, conducted the systematic review, and wrote the first draft of the manuscript. All authors subsequently contributed to improvement of the manuscript.

